# Curating clinically relevant transcripts for the interpretation of sequence variants

**DOI:** 10.1101/276287

**Authors:** Marina T. DiStefano, Sarah E. Hemphill, Brandon J. Cushman, Mark J. Bowser, Elizabeth Hynes, Andrew R. Grant, Rebecca K. Siegert, Andrea M. Oza, Michael A. Gonzalez, Sami S. Amr, Heidi L. Rehm, Ahmad N. Abou Tayoun

## Abstract

Variant interpretation depends on accurate annotations using biologically relevant transcripts. We have developed a systematic strategy for designating primary transcripts, and applied it to 109 hearing loss-associated genes that were divided into 3 categories. Category 1 genes (n=38) had a single transcript, Category 2 genes (n=32) had multiple transcripts, but a single transcript was sufficient to represent all exons, and Category 3 genes (n=38) had multiple transcripts with unique exons. Transcripts were curated with respect to gene expression reported in the literature and the Genotype-Tissue Expression Project. In addition, high frequency loss of function variants in the Genome Aggregation Database, and disease-causing variants in ClinVar and the Human Gene Mutation Database across the 109 genes were queried. These data were used to classify exons as "clinically relevant", "uncertain significance", or "clinically insignificant". Interestingly, 7% of all exons, containing >124 "clinically significant" variants, were of “uncertain significance”. Finally, we used exon-level next generation sequencing quality metrics generated at two clinical labs, and identified a total of 43 technically challenging exons in 20 different genes that had inadequate coverage and/or homology issues which might lead to false variant calls. We have demonstrated that transcript analysis plays a critical role in accurate clinical variant interpretation.

## Introduction

With the rapid growth of genomic testing and the dropping cost of sequencing, proper analysis of genetic variants is critical for patient care. The American College of Medical Genetics and Genomics (ACMG) has set forth guidelines for the interpretation of sequence variants ^1^. However, use of the guidelines requires an understanding of the transcriptional architecture of each gene. There can be several mRNA transcripts for each gene, and each laboratory individually determines which transcript to use when annotating, interpreting, and reporting variants in any gene. Human transcripts are currently designated and annotated by multiple groups. The most commonly used sets of coding transcripts that are currently available come from GENCODE (Ensembl and HAVANA, CCDS, LRG), UCSC, RefSeq (LRG), and AceView^2-8^.Each group annotates transcripts with a combination of computational and manual literature curation. Although current HGVS standards mention that describing variants in the context of exons and introns is optional ^9^, in many genes there are exon-specific factors that influence interpretation and tracking this data on an exon level is important.

In addition to the above annotation challenges, technical limitations of NGS can also lead to inaccurate variant calls. Several genes contain coding sequences that can pose several technical problems, including sequences with high homology to other genomic regions, high GC content, and repetitive sequences. If a gene has significant sequence overlap with another gene or a pseudogene, it can be difficult to align the short NGS reads to the right genomic location, leading to false negative and/or false positive variant calls. DNA with high GC content is not easily amplified, and highly repetitive DNA is prone to sequencing and/or alignment errors. All such regions should be systematically investigated in the targeted genes of interest to address test limitations and design necessary ancillary assays ^10^.

Upon passing sequencing quality metrics and filtration cutoffs, a rare variant would then be evaluated based on the most biologically relevant transcript for the disease of interest. It is common to choose the longest transcript for sequencing pipelines. However, variants are often evaluated in the context of this transcript, which does not necessarily encompass all essential exons and can also contain non-biologically functional exons. Thus, choosing a medically relevant transcript is essential for variant interpretation, and for understanding the molecular consequence of a variant on the gene’s function.

Here we provide a framework for transcript curation and selection using a combination of tissue expression and genomic datasets, protein functional domains, and published work from animal and human studies. We apply this framework to hearing loss, a relatively common condition that affects 1 in 300 infants, half of which have a genetic etiology ^11^. Due to the complexity of the auditory system, it is also highly heterogeneous with over 100 genes causative for nonsyndromic hearing loss alone ^12^. We also use clinical NGS datasets generated at two different diagnostic laboratories to systematically highlight technically challenging regions across the hearing loss genes. We demonstrate the utility of our framework and its impact on variant annotation and interpretation. While our analysis was limited to hearing loss, we recommend that this guidance be used for all genes that are definitively associated with fully penetrant diseases.

## Methods

### Transcript Curation process

109 hearing loss-associated genes largely from the OtoGenome™ Test (GTR000509148.8) at the Laboratory for Molecular Medicine (LMM) were included for transcript curation. All known (NM) RefSeq transcripts in these genes were curated with respect to function, tissue specificity and temporal expression from published literature (**Figure 1**). Exon-specific expression data were extracted from the Genotype-Tissue Expression Project (GTEx) on 01/15/18. GTEx was supported by the Common Fund of the Office of the Director of the National Institutes of Health, and by NCI, NHGRI, NHLBI, NIDA, NIMH, and NINDS. In addition, high allele frequency (>0.3%) predicted loss of function (LoF) variants (nonsense, frameshift and +/-1,2 splice site) were queried from the Genome Aggregation Database (gnomAD) ^13^, and exons containing such LoFs were flagged. An allele frequency of 0.3% was chosen since variants in hearing loss genes that are above this frequency can be considered likely benign as defined by Duzkale et al ^14^.

**Figure 1:**
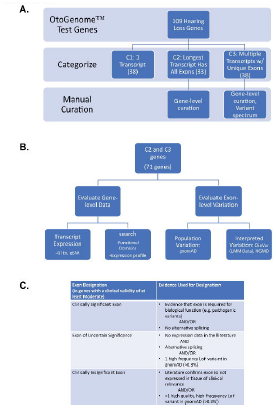
**A) Transcript Curation Workflow:** 109 hearing loss-associated genes, predominantly from the OtoGenome™ Test (GTR000509148.7), were categorized. Genes were divided into 3 categories using NCBI reference sequence (RefSeq) transcripts. Category 1 (C1) contained genes that had a single transcript, genes in category 2 (C2) had multiple transcripts, but the longest transcript encompassed all exons, and Category 3 (C3) genes had multiple transcripts with unique exons. **B) Category 2 and 3 Curation Process:** Category 2 and 3 genes were manually curated. Exon-specific expression data was pulled from GTEx and gEAR. Literature searches were performed for information about functional domains, additional expression data, such as tissue-specific transcript expression, and temporal expression. To evaluate population variation, loss of function variants were pulled from gnomAD. To evaluate interpreted variation, LP/P variants were pulled from our internal database (also in ClinVar) and ClinVar and DM variants were pulled from HGMD. **C) Exon Curation Process:** Exons were categorized as “Clinically relevant,” "Uncertain significance, “or “Clinically Insignificant” based on the pieces of evidence listed in the table.

### Exon Numbering and Classification

All coding and noncoding exons in the primary transcript were numbered sequentially. Coding and noncoding exons in minor transcripts were also numbered separately and sequentially per transcript. Transcripts were aligned and viewed using Alamut (Version 2.6.1 Interactive Biosoftware). Each minor transcript was given a different letter that was added after the exon number. For example, if there were two transcripts with unique alternate exon 1, they were numbered as 1A and 1B. To define a minimal curated transcript list, unique exons were listed in the minimal number of minor transcripts and the designated longest transcript contained the most coding bases **(Table 1, Supplemental Tables 1-3)**.

**Table 1:**
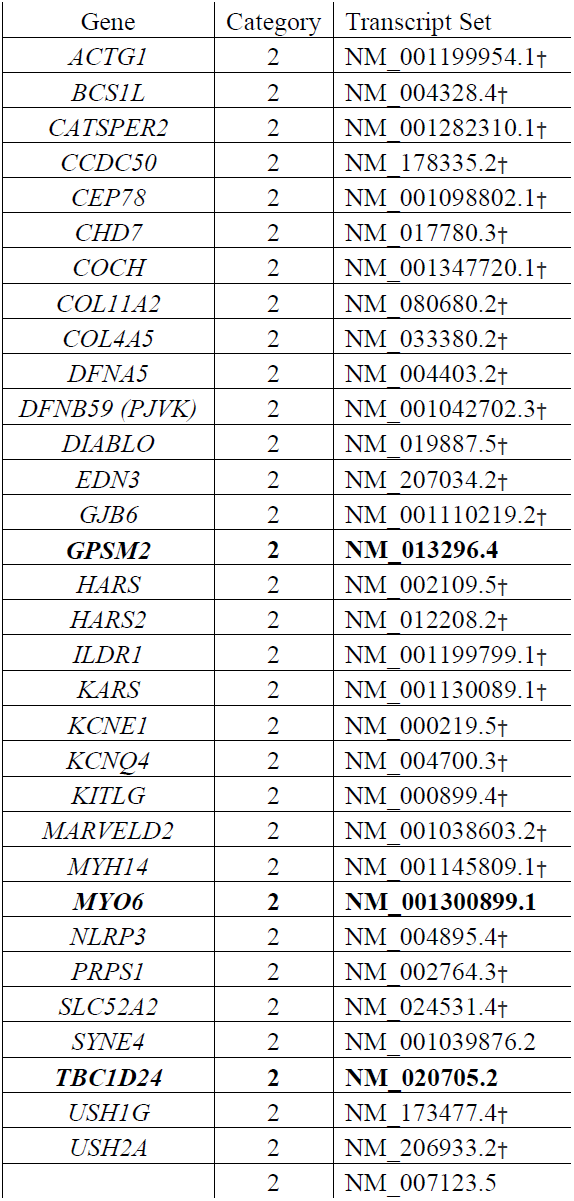
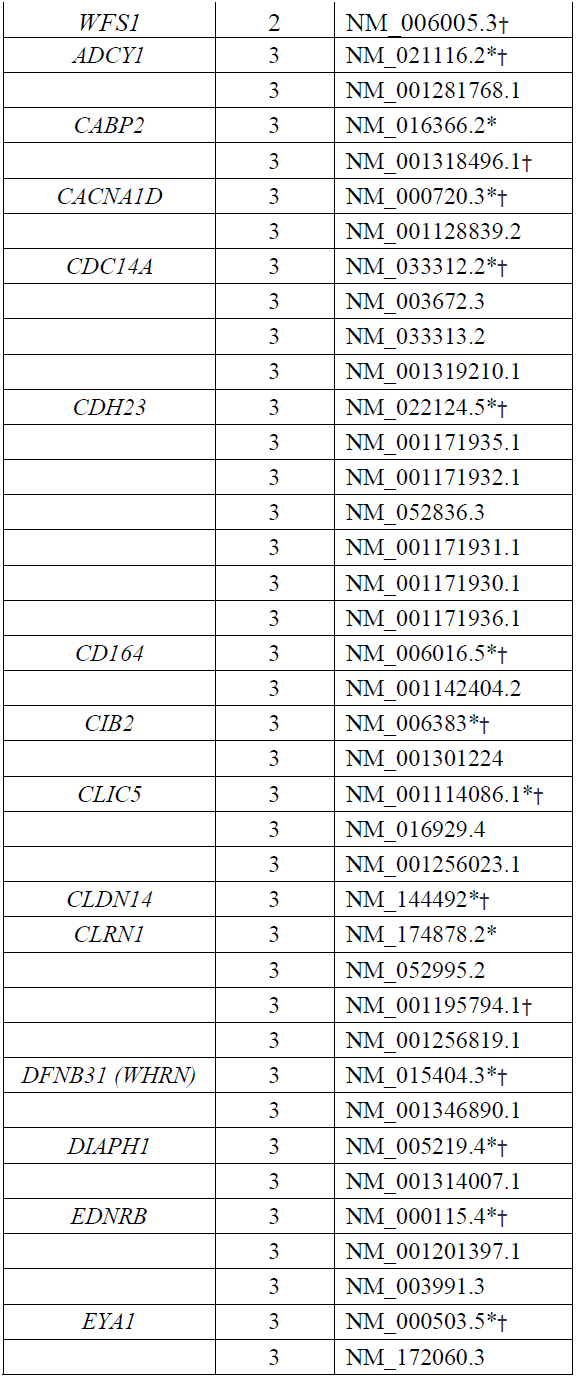
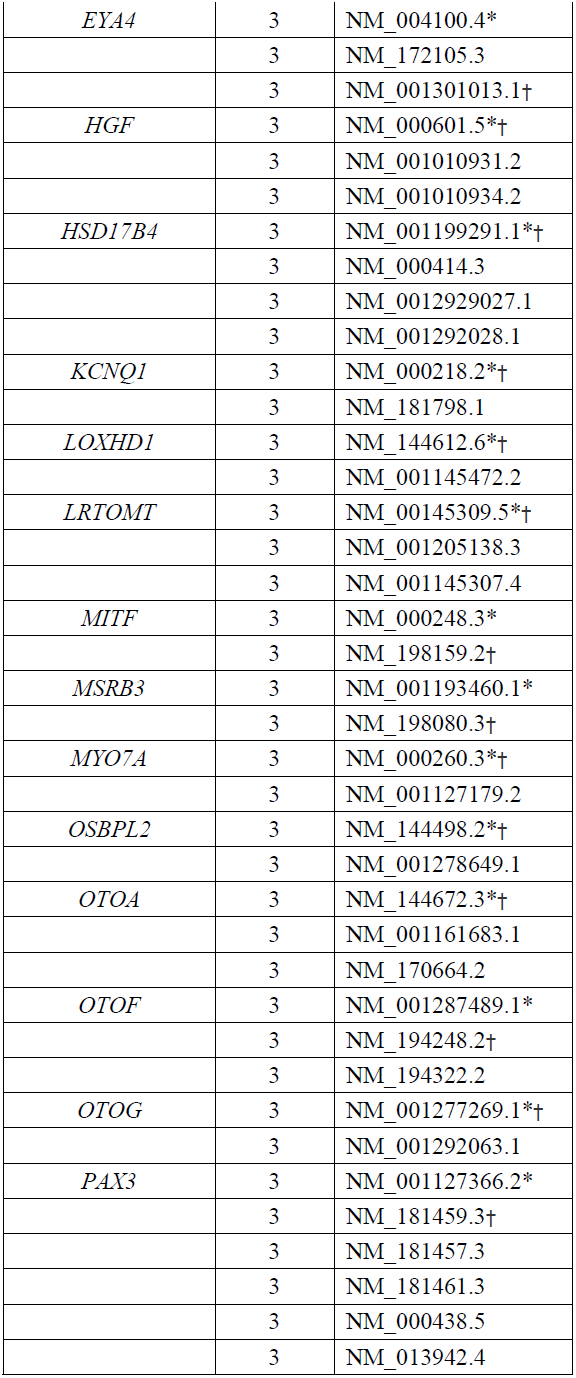
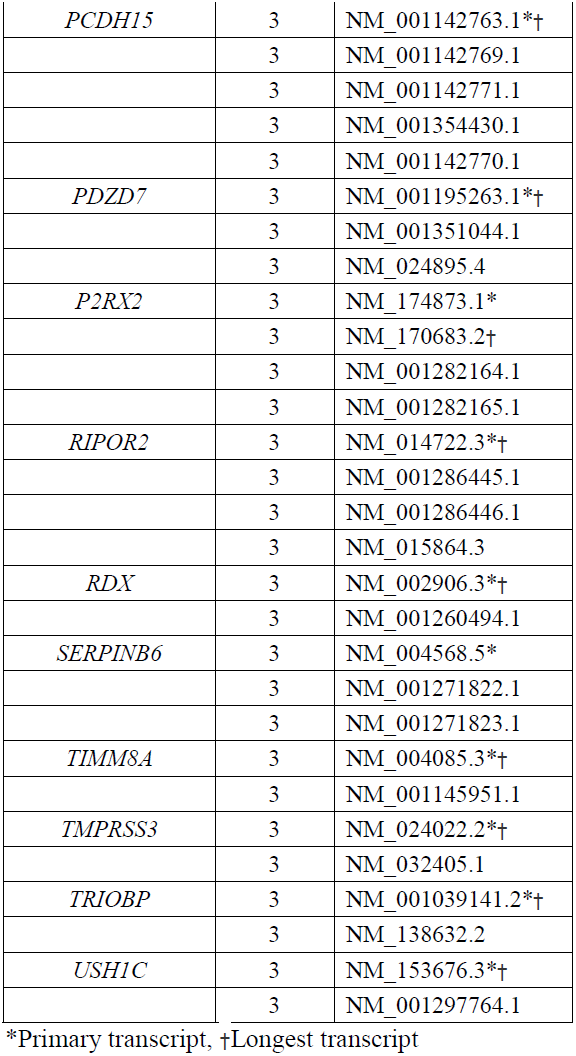
Curated transcripts for category 2 and 3 genes. The minimal curated transcript set with all unique exons for categories 2 and 3 are listed. C2 genes in which the curated transcript is not the longest one are listed in bold.

Exons were classified as "clinically significant", "uncertain significance", or "clinically insignificant" **(Figure 1C)**. Exons were classified as "clinically significant" if there was no evidence they were alternatively spliced, they did not contain high frequency exonic LoF variants, or they were supported by tissue-specific inner ear expression in the literature. "Uncertain significance" exons were spliced out of major transcripts, had no expression data and, for some, contained one high frequency LoF variant. Finally, "clinically insignificant" exons were noncoding, had non-supporting human or animal tissue expression data, or had multiple high frequency LoF variants.

### Variant Counts

Pathogenic (P), likely pathogenic (LP), benign (B), likely benign (LB) variants, and variants of uncertain clinical significance (VUS) in the ClinVar database ^15^ in addition to Disease Mutations (DMs) in the Human Gene Mutation Database (HGMD) ^16^ were counted across all uncertain and insignificant exons. Each variant was evaluated based on transcript location and predicted molecular consequence to the gene.

### Technically challenging regions

Exon-level next generation sequencing (NGS) quality metrics, including average mapping quality (MQ) and average minimum depth of coverage (DP), were calculated across all 109 genes using exome sequencing data. Exome targets were captured using the Agilent Clinical Research Exome (CRE) V5 kit, after which 2x100bp paired-end sequencing was performed on the Illumina HiSeq platform.

Exome sequencing using the above conditions was performed at the Children’s Hospital of Philadelphia (CHOP) ^17^ and the Laboratory for Molecular Medicine (LMM) with an overall average coverage of ~100x and 180x, respectively. Poor quality regions as defined in **Results** section were compared between the two sites. Additionally, 87 of the 109 genes were targeted at the LMM using a custom-based capture kit (Agilent), followed by 2x150bp paired-end sequencing to an average coverage of ~600x. Exon-level NGS quality metrics from this capture-based sequencing approach were calculated and compared to those from exome sequencing.

## Results

### Transcript curation

A total of 109 genes on LMM’s hearing loss panel were curated for clinically relevant transcripts as outlined in **Figure 1**. These genes had between 1 and 17 NCBI reference sequence RefSeq transcripts and between 1 and 72 unique exons, for a total of 340 unique transcripts and 2161 unique exons across all genes (**Figure 2A**). Genes were divided into 3 categories using RefSeq transcripts ^5^. All genes with only one RefSeq transcript were classified as category 1 (C1) genes. Genes with multiple transcripts were considered category 2 (C2) genes if the longest transcript included all annotated exons and those with multiple transcripts and mutually exclusive exons were grouped under category 3 (C3) (**Figure 1**).

**Figure 2:**
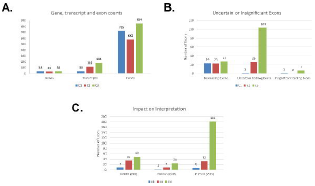
A) Gene, exon, and transcript Counts: Genes were categorized and transcripts and exons were counted. **B) Exon Counts Across Categories:** For each of the three categories, exons were classified as Noncoding, Uncertain, or Insignificant as per the definitions in Figure 1C. **C) Impact on Interpretation:** Variant counts in uncertain and insignificant exons for each category were collected from HGMD and ClinVar. DM, Disease causing mutation; LP, Likely Pathogenic; P, Pathogenic; VUS, Variant of Uncertain Significance.

#### Category 1 genes

Of the 109 genes evaluated, 38 had a single RefSeq transcript each with an average of 19 exons (total of 38 transcripts and 725 exons across 38 genes, **Figure 2A** and **Supplementary Table 1**). Because each gene had only one known RefSeq transcript, C1 transcripts had minimal curation and almost all exons, except for noncoding ones (n=24), were considered to be critical for inclusion in diagnostic testing and variant interpretation. Interestingly, however, based on exon-level counts of high allele frequency loss of function (LoF) variants in the general population (see **Methods**), there was one insignificant exon (*MYO15A*, NM_016239.3, Exon 26) and another of uncertain significance (*ATP6V1B1*, NM_001692.3, Exon 1) in this category.

The *MYO15A* exon 26 contained a nonsense variant (c.5925G>A; p.Trp1975*) that was present in 1.3% (316/23,712) South Asian alleles including 3 homozygotes in the Genome Aggregation Database (gnomAD). Interestingly, RNA sequencing data from the Genotype-Tissue Expression (GTEx) study showed that this exon is not expressed across all tested tissues. We therefore considered this exon to be clinically insignificant and that variants identified therein are more likely to be benign. The *ATP6V1B1* exon 1 contained a start loss variant (c.2T>C; p.M1?) that was present in 40% total alleles in gnomAD, including 23,280 homozygotes. It is possible that the exon start is erroneously annotated or that re-initiation might occur elsewhere, including at any of the two downstream methionines in this exon. However, there is currently no functional data to support either possibility or to rule out potential re-initiation at downstream exons. We therefore classified this exon as uncertain clinical significance wherein sequence variants should be carefully interpreted.

#### Category 2 genes

This category included 33 genes each with an average of 4 transcripts and 18 unique exons (**Figure 2A** and **Supplementary Table 2**). A total of 118 transcripts, including 582 unique exons, were curated in this category. Although the longest transcript represented all annotated RefSeq exons and was presumed to be the major transcript for C2 genes, we curated the shorter (minor) transcripts for potentially identifying non-biologically or non-clinically relevant exons in the longer transcript. Apart from the 23 noncoding exons in these genes, there were 26 coding exons not contained in minor transcripts, thus questioning their clinical relevance (**Figure 2B**). Of those, 7 exons were not expressed in any tissue in the GTEx database including 4 exons (*CCDC50* exon 6, *DFNA5* exon 2, *EDN3* exon 4, and *ILDR1* exon 6) harboring high allele frequency LoF variants in gnomAD (**Supplementary Table 2**). Of the 15 exons that showed expression in the GTEx database, 5 exons (*CEP78* exons 1, 2, and 16, *DFNA5* exon 6, and *KARS* exon 15) also contained high frequency LoF variants in gnomAD (**Supplementary Table 2**), while the remaining 10 exons did not, although more information is needed to clarify their biological or clinical relevance.

An illustrative example in this category is the *EDN3* gene known to cause Waardenburg syndrome (WS) type 4 ^18 19^. This gene has 5 RefSeq transcripts sharing coding exons 1-3 and 5 but differing in the inclusion of coding exon 4. Specifically, the NM_001302456.1 and NM_207033.2 transcripts do not contain the fourth coding exon shared by the three other transcripts (**Figure 3**). Interestingly, a frameshift variant (NM_207034.2: c.559_560insA; p.Thr189Asnfs) in this exon was present in 0.6% (157/25790) Finnish European alleles in gnomAD, with a high quality variant score, including 2 homozygotes (**Figure 3**), supporting the clinical insignificance of this exon.

**Figure 3:**
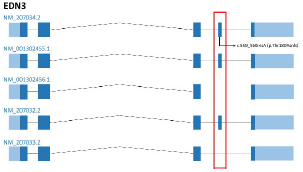
Visualization of category 2 example, *EDN3*. **A)** Transcript view of the *EDN3* gene. The high frequency loss of function variant is pulled from gnomAD and is located in the exon boxed in red.

#### Category 3 genes

There were 38 genes in this category, each with an average of 5 transcripts and 22 unique exons (**Figure 2A** and **Supplementary Table 3**). In total, C3 genes had 184 RefSeq transcripts and 854 unique exons. Given the multiple transcripts with mutually exclusive exons in this category, a thorough curation was carried out to select the most clinically relevant transcript for each C3 gene (**Supplementary Table 3**). Published human and/or animal tissue expression studies supported transcript selection for 30 C3 genes; the longest transcript was supported in 25 genes while a shorter isoform was most relevant in 5 genes. There were no expression data to guide selection of the most biologically relevant transcript for 8 C3 genes for which we defaulted to the longest transcript. Overall, 104 coding exons in the C3 genes met our criteria for “uncertain significance“, while 7 coding exons were classified as "clinically insignificant" (see **Methods** and **Figure 2B**).

A C3 example is the *PAX3* gene which is a common cause of WS, type 1 ^20-22^. This gene has 8 RefSeq transcripts with varied tissue and temporal expression ^23-25^, and with significant alternative splicing; a transcript can include 4, 5, 8, 9, or 10 exons. Certain exons use alternate splice junctions which can also change reading frame for the terminal exon (**Figure 4A**). Interestingly, one putative LoF variant (c.638C>A; p.S213*) in exon 4d of the NM_000438.5 transcript was found in 0.54% (166/30,592) South Asian alleles including one homozygous individual. This allele frequency is inconsistent with the estimated disease prevalence for autosomal dominant WS of approximately 1/40,000^22,26^. Exon 4d is only 20 amino acids longer than exon 4 on all other transcripts. Human expression data in the GTEx database strongly supports usage of the exon 4 (and not 4d) splice donor site, suggesting that exon 4d is not biologically relevant (**Figure 4B**).

**Figure 4:**
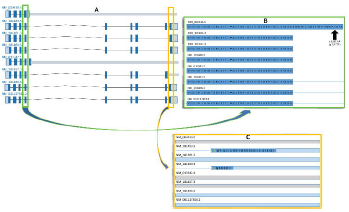
Visualization of category 3 example, *PAX3*. **A)** Transcript view of the *PAX3* gene. **B)** A close-up of the high frequency nonsense variant in exon 4d. **C)** A close-up of the two uncertain exons in PAX3, and 9a (NM_181459.3) and 8c (NM_181460.3).

Similarly, exon 8c in the NM_181460.3 transcript of the *PAX3* gene is unlikely to be biologically relevant as supported by lack of its expression (GTEx database) and the presence of a putative LoF splicing variant (c.1024+1G>C) impacting this exon in 0.15% (47/30,764) South Asian alleles in gnomAD. This same variant has a missense effect (p.Arg402Pro) on the NM_181461.3 transcript, further highlighting the importance of appropriate transcript selection for variant annotation and interpretation **(Figure 4C)**.

### Impact on clinical testing and interpretation

Transcript selection can significantly alter variant interpretation because variants can have differing molecular consequences on each transcript. For example, pathogenic missense and nonsense variants in the *MITF* gene, encoding a transcription factor critical for melanocyte development, are known to cause WS, type 2 ^22,27^. This gene has 13 curated RefSeq transcripts. On 4 out of 13 transcripts, a particular pathogenic variant (which segregated with disease in >10 members of a family with WS ^27^) is annotated as a variant in a +1 canonical splice donor site (c.33+1G>A). However, this nucleotide change is a deep intronic variant in the other 9 transcripts (e.g. NM_001184967.1:c.199-1066G>A), and could easily be misclassified if only the deep intronic consequence were interpreted.

Tissue-specific transcript expression can also significantly alter variant interpretation. Variants in *TBC1D24*, a GTPase-activating protein, are associated with either nonsyndromic hearing loss, DOORS syndrome, or a spectrum of epilepsy conditions ^28-33^. This gene has 2 curated RefSeq transcripts: NM_001199107.1, the longest which contains 8 exons and is most abundant in mouse neurons, and NM_020705.2, which is missing exon 3, contains only 7 exons, and is expressed in mouse cochlea and non-neuronal tissues ^29^. NM_001199107.1:c.969_970delGT (p.Ser324Thrfs), a frameshift variant in exon 3, is not present in the shorter transcript and was identified in the homozygous state in five members of a consanguineous family with severe lethal epileptic encephalopathy but no hearing loss, thereby supporting the tissue-specific expression of the longest transcript ^31^.

Based on our comprehensive curation of all transcripts, there were 139 coding exons with no or uncertain clinical significance, constituting 7% of all 2089 coding exons across all 109 genes (**Figure 2B**). Because of the limited evidence supporting those exons’ clinical relevance, variants therein should be carefully interpreted as they can be a source of false positive diagnoses. Interestingly, there are 124 variants that are labeled as disease causing (DM or P/LP) in disease databases (HGMD and ClinVar, respectively), in addition to 224 VUSs and 151 B/LB variants across those exons (**Figure 2C**). These variants interpreted as clinically significant will all require further assessment to ensure sufficient evidence is present to implicate them in hearing loss. This highlights the importance of our transcript curation approach and the impact it could have on functional and clinical annotations.

### Technical Assessment

Due to its genetic heterogeneity, most clinical genetics laboratories use targeted or exome-based panels to sequence a comprehensive set of genes known to cause hearing loss. Although very robust, such approaches have limitations inherent to the next generation sequencing (NGS) technology including the inability to reliably capture and sequence low complexity and/or high homology genomic regions.

We sought to identify regions in a set of 109 hearing loss genes that are technically challenging to sequence using current short read (100-150bp) NGS in clinical laboratories. We used clinical exome sequencing data generated in two different sites (CHOP and LMM) to calculate exon-level quality metrics across all exons in the 109 hearing loss genes. We have recently shown that an average mapping quality (MQ) and/or an average minimum depth of coverage (DP) cutoffs of 20 and/or 15, respectively, are strong indicators of poor quality regions ^17^. We identified 43 well-baited (~90% baited bases) exons in 20 genes with the above cutoffs despite being exome sequenced to an overall average coverage of up to 180x at the two clinical laboratories (**Supplementary Table 4** and **methods**). Of those, 31 exons were sequenced to an overall coverage of ~600x using a different targeted capture (average % baited exons: 95%) and longer reads (150bp), but still had low quality metrics (**Supplementary Table 4**).

The 43 regions included exons with high homology to other genomic sequences (n=21 exons in *STRC* and *OTOA*) or exons that have GC-rich or repeat sequences (n=22, e.g. *KCNQ1, MYO15A*, and *TPRN*) (**Figure 5**). It is unlikely that sequence variants in all 43 exons will be reliably detected using available NGS chemistries, and therefore false positive and/or false negative variant calls in those exons should be highly expected.

**Figure 5.**
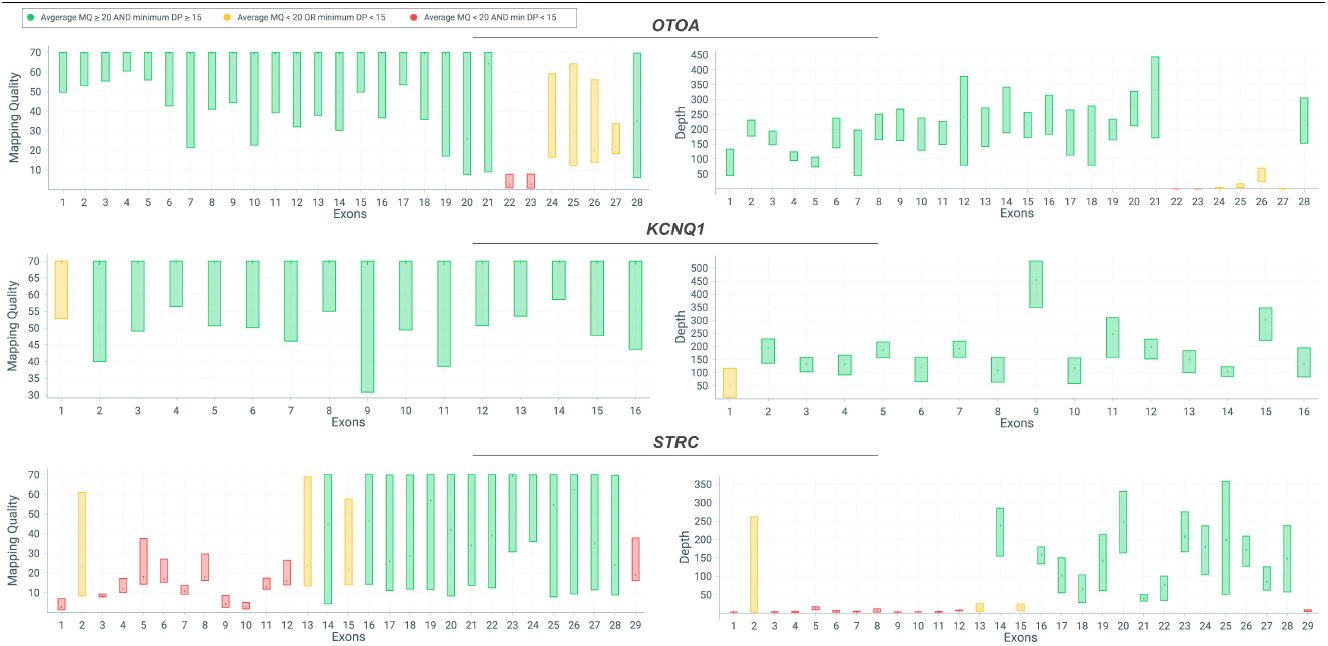
Visualization of three genes with known technically challenging regions. Exons in Otoancorin, *OTOA* (*top*) or Stereoclin, *STRC* (*bottom*) with high homology to other genomic sequences; and GC-rich first exon in the potassium channel, *KCNQ1* (*middle*). MQ and coverage plots are displayed for *OTOA, KCNQ1* and *STRC*. Green bars indicate exons with both average MQ ≥ 20 and min DP ≥ 15. Orange bar represents either average MQ < 20 or min DP < 15. Red bars indicate poor exons where both average MQ < 20 and min DP < 15. Each bar in MQ and coverage plots shows minimum and maximum range for each exon (top and bottom of the bar), average is shown by a tick mark in the middle of each bar.

## Discussion

Transcript selection is critical for determining DNA variants’ potential effects on RNA and/or protein expression, function and stability. This annotation, in turn, significantly impacts variant interpretation. Although most clinical laboratories use one set of coding transcripts (commonly RefSeq) for variant annotation, any set might contain multiple transcripts for each gene; some of which are true clinically relevant isoforms, while others can be false annotations. Even for those genes with multiple isoforms, deciphering the relevant isoform for a given disease – often based on tissue-specific expression data – is necessary for interpretation. In the absence of uniform guidelines for transcript selection, each lab applies different internal rules for identifying the most appropriate transcript(s) for interpretation and reporting.

Here we provide a comprehensive evidence-based framework for transcript curation and selection. We apply this framework to 109 hearing loss genes, and illustrate its utility in transcript selection and variant annotation and interpretation. We also use a new exon classification system, and show that 7% of all coding (RefSeq) exons in these genes have no or questionable clinical validity rendering them a potential source of false variant calls irrespective of their predicted protein effect (missense, loss of function, etc.).

A challenge with our approach is that it requires significant manual curation, though such curation is essential for accurate interpretation, and is arguably more effective if performed ahead of testing, and not retrospectively to minimize analysis and wet bench burdens. Another challenge is that it is highly dependent on availability of human and/or animal expression data in the relevant disease tissue – the inner ear in this current work. However, leveraging existing large human genomic population (gnomAD) and transcriptome (GTEx) sequencing data as well as high quality variant databases (ClinVar) can support the selection of clinically relevant transcripts in our genes.

Finally, we use exon-level NGS quality metrics to highlight regions that are inaccessible to sequencing and/or accurate variant calling, especially with short read (100-150bp) chemistries that are mostly used in clinical and research labs. It is possible that some of those regions can be recovered with longer reads and improved bioinformatics pipelines. Until then, however, it is highly important that different ancillary assays, such as Sanger sequencing, be validated to accurately capture sequence variants in those regions.

In summary, we recommend that our transcript selection framework and exon classification system be used in other disease areas for more efficient and accurate variant interpretation, and to avoid erroneous annotations and, potentially, misdiagnoses.

## Acknowledgements

Research reported in this publication was supported by the National Human Genome Research Institute (NHGRI), in conjunction with additional funding from the Eunice Kennedy Shriver National Institute of Child Health and Human Development (NICHD) under award number: U41HG006834. The content is solely the responsibility of the authors and does not necessarily represent the official views of the National Institutes of Health.

